# Temporal and spatial patterns of vertebrate extinctions during the Anthropocene

**DOI:** 10.1101/2022.05.05.490605

**Authors:** Daniel Pincheira-Donoso, Lilly P. Harvey, Jacinta Guirguis, Luke E. B. Goodyear, Catherine Finn, Jack V. Johnson, Florencia Grattarola

**Author notes:** Corresponding author: D. Pincheira-Donoso.

## Abstract

The human-induced annihilation of modern biodiversity is dragging the planet into a mass extinction that has already altered patterns of life globally. Among vertebrates, over 500 species have become extinct or possibly extinct in the last five centuries – an extinction rate that would have taken several millennia without human intervention. Vertebrate extinctions have often been quantified as cumulative counts that reveal sharp increases in losses over time. Here, we quantify global tetrapod extinctions since the 1400s using numbers of species losses across successive and independent time periods until present. Our results reveal that extinctions were low and fundamentally restricted to islands in pre-industrial times, experiencing a significant increase and spread over continental mainland following the onset of the industrial revolution. Recent amphibian extinctions alarmingly exceed the extinctions of all tetrapods, while extinctions of island birds account for a third of all extinctions. Finally, we quantified the relationship between human population growth (HPG, as a proxy for aggregate human effects on the environment) and extinctions between 1800-2000, to then predict that an estimated 838 tetrapod species will go extinct between 2030-2100 based on United Nations HPG projections. These findings further warn humanity about the need to sustainably control HPG and the destructive impacts of rapid environmental change on ecosystems worldwide.

## Introduction

The rapid extinctions of global biodiversity rank among the most disturbing consequences of modern alterations of the planet’s environments caused by humans. The accumulation of evidence for the persistent loss of local populations and of entire species from across the tree of life^1,2^ has reinforced the warning that the modern era – the Anthropocene – is on the brink of entering the sixth mass extinction recorded on Earth^2–9^. The figures are staggering. Existing estimates suggest that modern rates of species extinctions range between 1,000-10,000 times higher than ‘background’ rates of extinction driven by ‘natural’ causes^10,11^, totalling the loss of nearly 1% of species, and several millions of populations only within the last few centuries^1,2,6,7,10,12–16^. Even more disturbingly, these figures are not only likely to be underestimated^10^, but also, the actual number of extinct species is expected to be drastically higher considering that around 60%-82% of animal species^10,17–19^ and at least 15% of plant species^10,20–22^ remain to be discovered. Finally, excluding already extinct species, more than a quarter of the world’s species whose conservation status has been assessed are threatened with extinction^23^. Collectively, this modern global extinction crisis is a rapidly expanding threat for the persistence of ecosystems globally, and for the welfare of life as a whole, including humans^2,6,24–26^.

In recent decades, a stream of global-scale analyses primarily conducted on vertebrates^6,7,15,27–31^ – guided by the International Union for Conservation of Nature (IUCN) Red List^23^ – have driven considerable progress in determining the quantitative patterns of modern species extinctions through geographic space and across lineages^1,2,6,7^, and the synergies among threats and population features that underlie these patterns^1,2,15,28,29,32–35^. Evidence reveals that the geographic and taxonomic distribution of extinctions is heterogeneous, with some regions of the world (e.g., tropical America, Southeast Asia)^1,7,31^ and some lineages (e.g., amphibians)^10,15,36^ concentrating exceptionally more severe extinction levels than others. In contrast, factors such as small geographic range size, habitat loss and low fecundity tend to be consistently identified as drivers of extinctions regardless of region or lineage^10,15,33,35^.

Combined, patterns and processes of vertebrate species extinctions (and from across nature more widely^1,2,12,14,35,37,38^) demonstrate with alarming consistency the role of humans in the rapidly unfolding modern mass extinction. The magnitude of this extinction crisis has been quantified by recent studies that have reconstructed the cumulative trajectories (i.e., each extinction event added to previously counted extinctions) of vertebrate extinctions worldwide over the last few centuries since pre-industrial times to the present^6,10,39^. Extinction rates, relative to background (‘natural’) extinction rates, have drastically increased, with sharp accelerations taking place in post-industrial times from the 1800s^6^. However, less attention has been given to the global-scale fluctuations (rather than the cumulative trajectories) of vertebrate species extinctions across independent time periods. Therefore, whereas the constant accumulation of extinct species is well-established, a full account about how extinctions have fluctuated through successive and independent time periods, through geographic space, and across vertebrate classes remains unclear. Here, we present the first analyses of the spatial patterns of the world’s tetrapod species extinctions through time since the end of the Middle Age when the first precise records of tetrapod extinctions (a mammal) in recent history are known^23^. We investigate (*i*) the temporal trajectories of extinctions recorded for each of the four tetrapod classes (amphibians, reptiles, birds and mammals), (*ii*) how the tendencies of extinctions have varied geographically and taxonomically over time, and (*iii*) predict future species extinctions as a function of human population growth – a paramount outcome of industrialization and the underlying cause of aggregate effects behind accelerating environmental degradation^40^.

## Results

### Temporal trajectories of extinctions

Tetrapod extinctions have increased over the past five centuries, with a significant acceleration for all taxa following the onset of the industrial revolution since post-1760 (Chi-squared, *X*^2^=15.837, *df*=1, *P*<0.001, Fig. 1a-e; Supplementary Table S1). Numbers of species extinctions on islands (*X*^2^=7.318, *df*=1, *P*=0.007) and on continents (*X*^2^=8.646, *df*=1, *P*=0.003) separately, have significantly increased in post-industrial times. This sharp increase is represented by the drastic change of extinction numbers shown in the curve for tetrapods combined (Fig. 1e). The earliest recorded extinctions since post-Middle Age (in the 1400s and 1500s) contain only birds and mammals (Fig. 1c-e), whereas the first records of ectotherms are known from the 1600s, when three species of reptiles (all squamates) became extinct (Fig. 1b; Supplementary Table S1). Until the onset of the industrial revolution period, the numbers of extinctions recorded every two decades for these three classes remained roughly constant (Fig. 1b-d; Supplementary Table S2), with no trend in extinctions on continents (Kendall’s *τ*=-0.053, *P*=0.856) or for all mammals (*τ*=-0.135, *P*=0.571), and only island birds showing an increase in extinctions (*τ*=0.565, *P*=0.005). The first records of amphibian species extinctions are known from the second half of the 1800s only (Fig. 1a). However, in contrast with the three other classes, amphibian extinctions have undergone an alarming acceleration, resulting in 122 species losses during the second half of the 1900s, which equates to over 140% of all other tetrapod extinctions combined during the same period (Fig. 1a, e, g). Even in the last 25 years, significantly more amphibian species have been lost than in the previous 25 years (*X*^2^=7.273, *df*=1, *P*=0.007), with a mean of two species lost every five years in 1951-1975, compared to 19 in 1976-2000. A similar situation is observed in extinctions that occurred after the year 2000, where the 22 species of extinct amphibians is nearly double the extinctions of all other tetrapods combined (13 in total; Fig. 1f; Supplementary Table S1). Evaluating numbers of extinctions post-1760, numbers of extinctions for all combinations of taxa and continent/island have been significantly increasing, apart from insular birds and insular amphibians, which are marginally significant (Supplementary Table S2).

**Figure 1.**
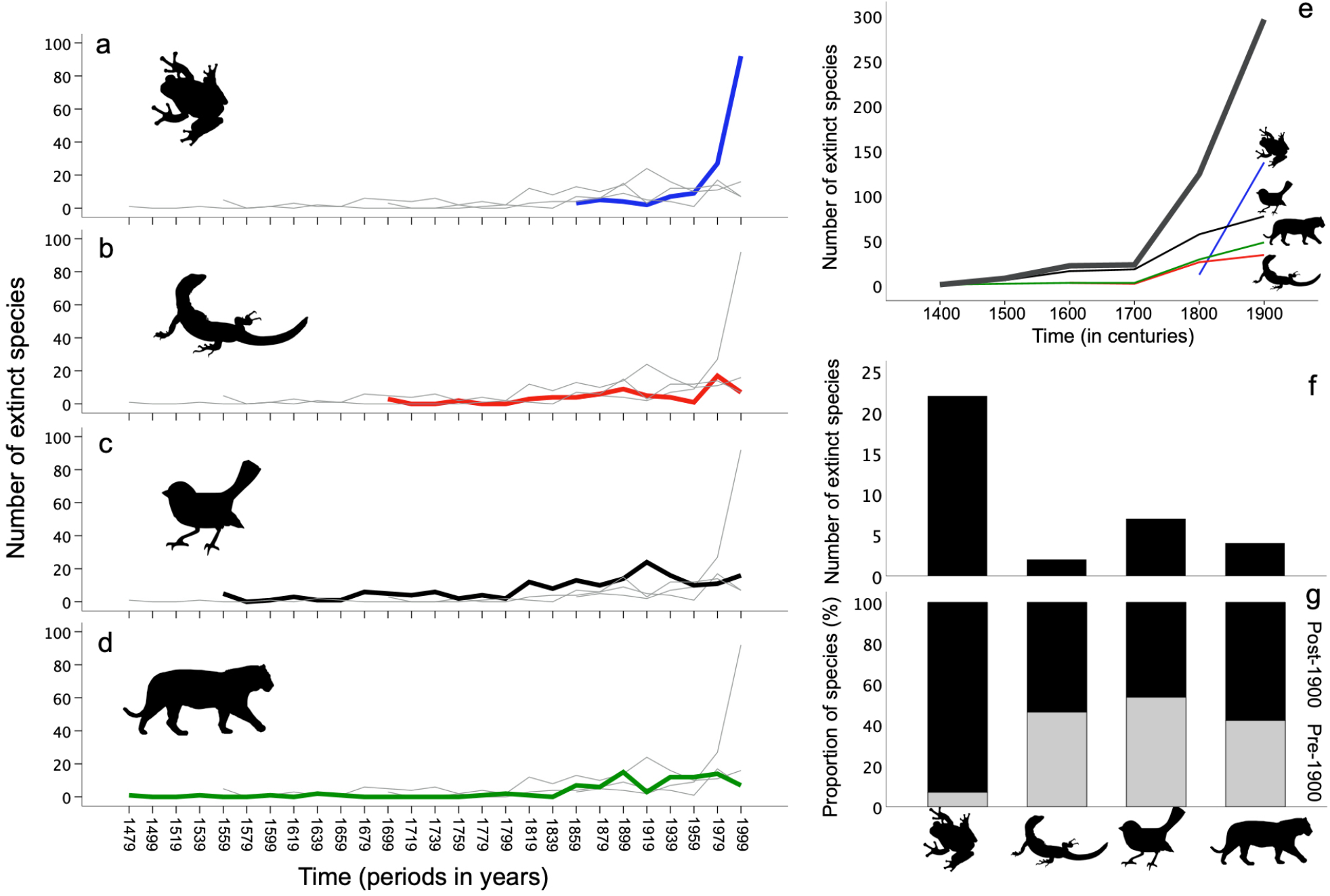
Temporal fluctuations in episodes of vertebrate species extinctions since the 15^th^ century. Extinction episodes plotted every two decades since the end of the Middle Age reveal a sharp increase in amphibian extinctions since the 1960s (a), in strong contrast with the slight, and fluctuating, overall increases in extinctions in reptiles (b), birds (c) and mammals (d). Extinctions in both classes of endotherms have much earlier records (c, d) than the observations in ectotherms (a, b). Temporal extinction trajectories of tetrapod classes separately across 100-year intervals, and combined (thick gray line), are shown in e. The numbers of species per class recorded to be extinct or possibly extinct since the year 2000 are shown in f, whereas the recorded year of extinction (or possible extinction) of species per class are shown separately for the periods before 1900 and after 1900 (g).

### Spatio-temporal patterns of extinctions

Tetrapod extinctions show a consistent biogeographic pattern through time, with species losses predominantly restricted to island endemics in pre-industrial times, followed by wide-spreading extinctions across continental mainland after the onset of the industrial revolution (Fig. 2; Supplementary Table S1). In the pre-industrial era, records of mainland extinctions only include two species (22.3% of extinctions recorded for the period) during the 1400s-1500s (one of which was in New Zealand), and four species (18.2%) during the 1600s (two of these in Madagascar, one in New Zealand), with no extinctions during the 1700s (Fig. 2; Supplementary Table S1). Overall, only 11.3% (five out of a total of 53 species) of all pre-1800s extinctions occurred on mainland (four of which occurred on >150,000km^2^ islands; see methods), whereas 88.7% of extinct species (48 species) were island endemics (Fig. 2). In stark contrast, 21 extinctions (16.9%) during the 1800s, and 171 extinctions (58.6%) during the 1900s occurred on mainland (Fig. 2). Overall, 46.2% (192 out of a total of 416 species) of all post-1800s extinctions took place on mainland, whereas 53.8% of species (224 species) were island endemics (Fig. 2). In fact, although most of these post-industrial mainland species extinctions are amphibians (123 species, 64% of all mainland extinctions), it is only from the 1800s that extinctions across all four tetrapod classes have consistently taken place on mainland. Throughout the entire historical period covered, island birds have been by far the most impacted of all tetrapods (Fig. 2), concentrating 58.1% (158 species) of all island endemic extinctions, and 33.7% of all island and mainland extinctions combined.

**Figure 2.**
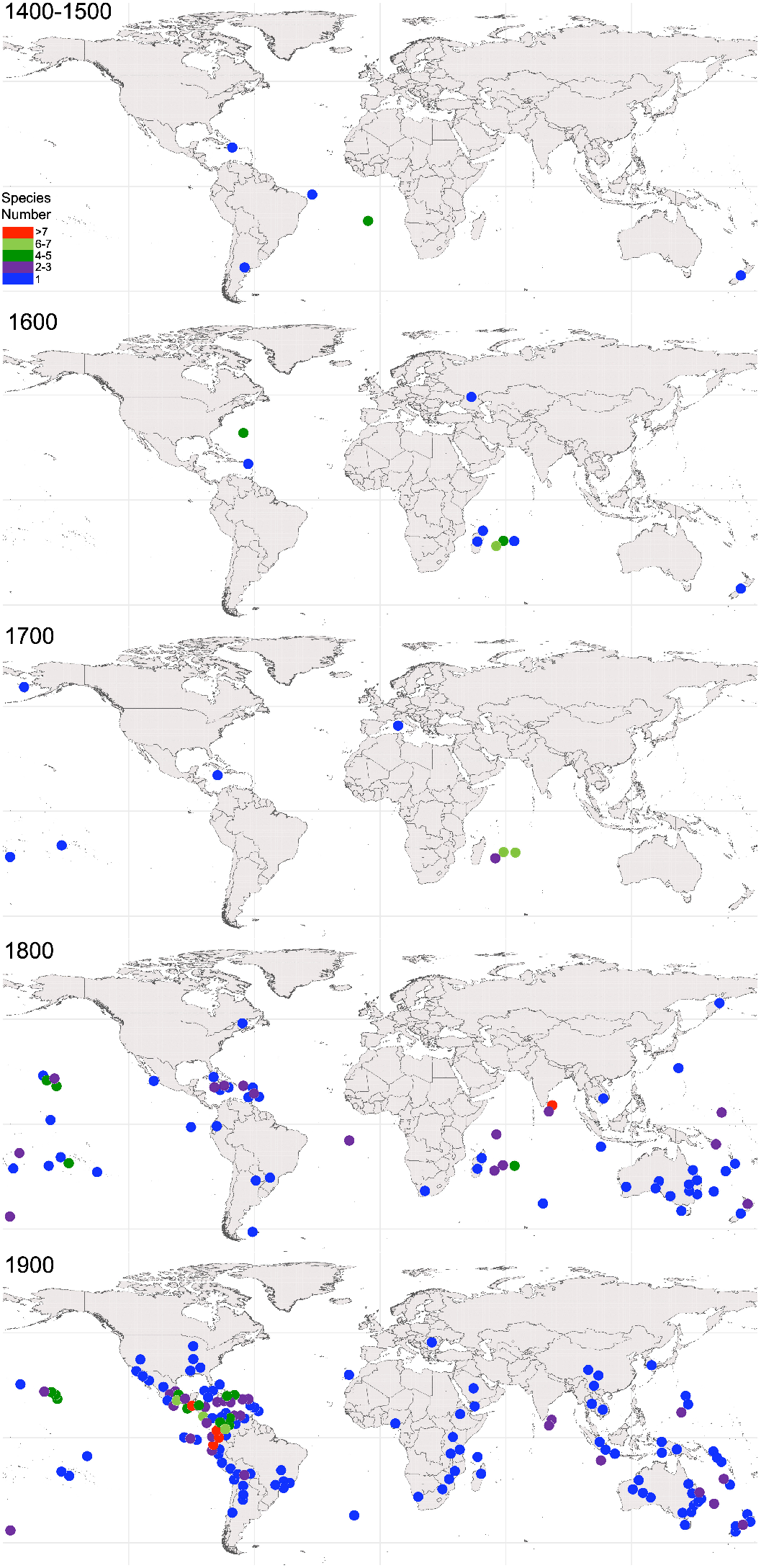
Spatiotemporal distribution of extinctions (and possible extinctions) of the world’s vertebrates since the Middle Age. There is a strong tendency for species extinctions to have concentrated on islands before the onset of the Industrial Revolution (1400-1799), in contrast with a drastic spread of extinctions over continental territories following industrialization (from 1800 onwards). Colour dots indicate 1×1 degree grid-cells in which the numbers of extinct species are 1-2 (blue), 3-6 (green) and 7-9 (red).

### Projections of future species extinctions

Predictions of tetrapod extinctions derived from the observed relationship between species losses and increasing human population growth (HPG) over time (quantified between 1800-2000) show a sharp increase of extinctions during the 21^st^ century (Fig. 3; Supplementary Table S3). Over the 2030-2100 period, using projections of human population growth data from the UN (see methods), an average of 55.87 species (range: 47 species between 2026-2030, 61 species between 2096-2100) are predicted to become extinct in every 5-year interval. In total, this results in an estimated 838 species predicted to be lost over that 70-year period (Fig. 3; see Supplementary Table S3, and Supplementary Figure S1), a figure equivalent to 155% of all tetrapod extinctions recorded since the 1400s.

**Figure 3.**
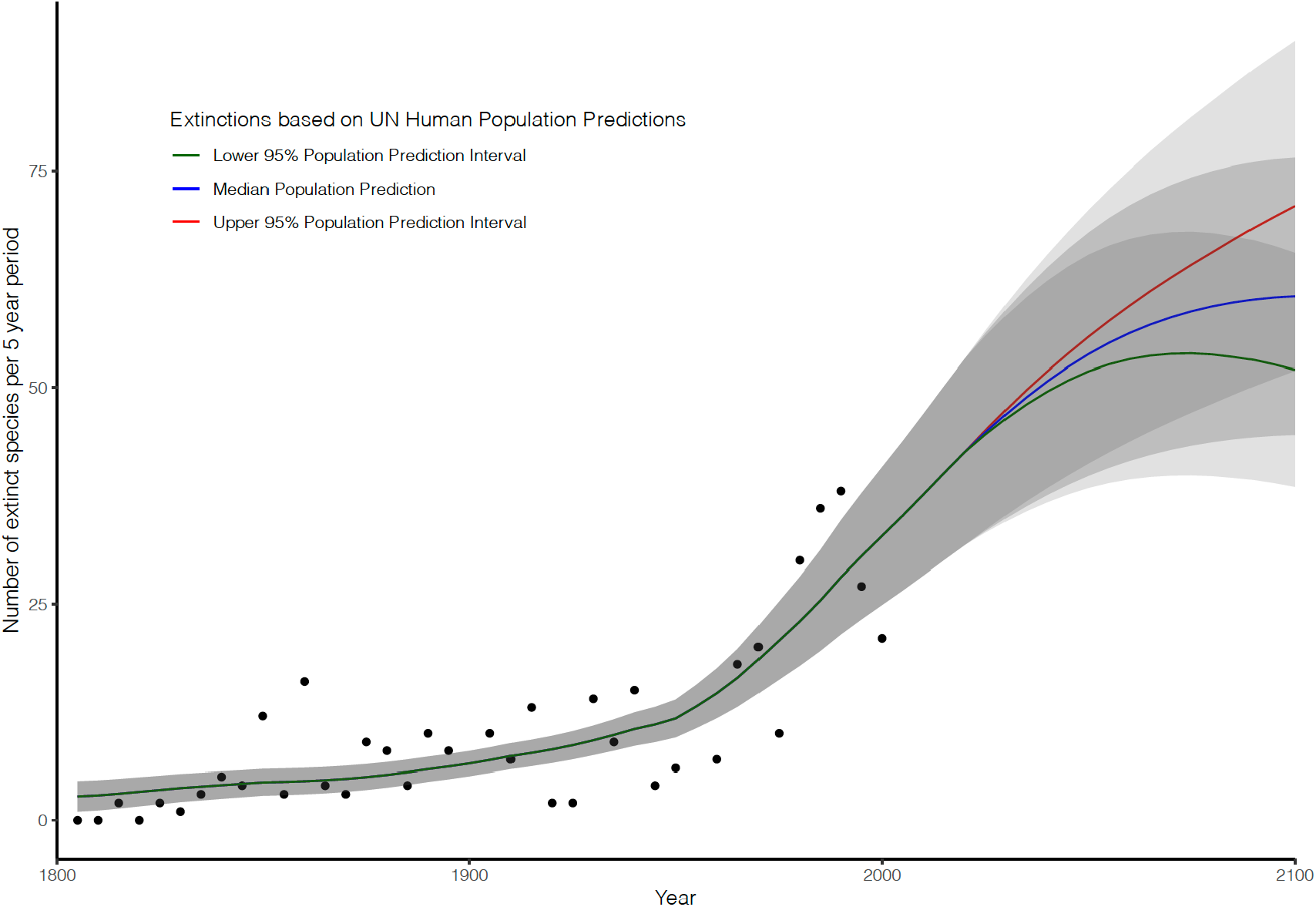
Number of recorded extinctions in 5-year intervals from 1801-2000 with predictions of number of extinct species in 5-year intervals between 2001 and 2100 based on human population growth scenarios from the UN. Green indicates the lowest bound of human population growth prediction, blue the median and red the upper bound. The numbers of extinctions are raw figures that indicate the number of species that are lost for every five-year period, with the year of each datapoint signifying the final year in the period. 95% confidence intervals are shown in grey.

## Discussion

Our results provide an in-depth description of the global spatio-temporal patterns of tetrapod species extinctions at successive periods of time over the past >550 years, since the end of the Middle Age, and projections of expected extinctions during the rest of the 21^st^ century. Unlike extinctions quantified as cumulative curves over time^6,10^, our analyses based on extinction counts across consecutive and independent time periods offer the advantage to better identify historical oscillations in species loss which can be linked to historical transitions in human activities. Two dominant extinction patterns are revealed by our study. First, that the numbers of species extinctions are relatively similar across tetrapod classes, with very low levels of extinctions in pre-industrial age, followed by significant accelerations in total number of species extinctions with the onset of the industrial revolution from the late 1700s^6^. Yet, some singularities exist. Mammal extinctions tended to remain particularly flat during pre-industrial age (with fluctuating post-industrial extinctions of mammals, birds and reptiles), whereas amphibian extinctions have increased explosively during the last century, with no extinctions recorded before the late 1800s. Although these tendencies might be at least partially influenced by paucity of records in pre-industrial age (e.g., see ref^41^), the overall extinction patterns shown by records are likely to accurately represent the actual trajectories of tetrapod extinctions over time. Second, whereas pre-industrial extinctions concentrated nearly entirely on islands, extinctions spread widely across continental landmasses only in the post-industrial era. Therefore, these patterns confirm the unambiguous role that anthropogenic environmental alterations caused by industrialisation (e.g., habitat destruction, rapid climate change) have played as causative agents of biodiversity extinctions at global scale.

Prior to the industrial era – when the scale of human population density^40^, industrial production, conversion of natural ecosystems into agricultural land^42,43^, CO_2_ emissions^40^, and exploitation of natural resources remained constantly low^1,44,45^ – the impacts of habitat loss and climate change on biodiversity were negligible. However, the introduction of invasive species was already a prevalent threat to biodiversity^41,46,47^, especially on islands, where the impacts of alien species are generally more severe given higher endemism and smaller geographic range sizes of native resident species^23,48–55^. In fact, a wide array of birds and mammals (as well as plants) were actively introduced all over the planet by European explorers during 1500s-1800s^23,49,50^, enabling the establishment of alien species worldwide. It is precisely some of these vertebrates that have been identified as prevalent causes of extinctions of island endemics over the past five centuries^52,56,57^, with rats (*Rattus*) being a prominent case of impact on biodiversity globally^55,58–60^. An additionally widely debated hypothesis suggests that drastic anthropogenic overharvesting (e.g., overhunting) might have been the cause behind rapid demographic collapses that led to extinctions of a range of species^61–64^. This explanation holds even for contested cases, such as the extinction of the Steller’s sea cow (*Hydrodamalis gigas*). This species is believed to have directly collapsed by excessive overhunting^62^, or indirectly by overhunting that caused the extirpation of the sea otter (*Enhydra lutris*) local populations, which had a keystone role in maintaining the shallow-water kelp forests on which sea cows fed^65,66^. Therefore, these scenarios are strongly consistent with our observation that the numbers of species extinctions in the pre-industrial era concentrated on islands.

The onset of the industrial age brought an unprecedented level of environmental and climatic alterations at global scale^1,43,44,67–69^, which not only added novel threats, but also aggravated the magnitude of human-induced effects that were already threatening biodiversity. For example, industrialisation facilitated a drastic increase in the spread of invasive species that were already driving island species to decline^41,47,49,70^. These changes also enabled the outbreak and spatial spread of infectious diseases with destructive impacts on wildlife^39,71–74^. The synergistic effects of these factors combined amplified the scale of anthropogenic impacts on ecosystems^35^ across both islands and mainland from the late 1700s^44^, thus expanding the threats on biodiversity and the rapid declines and extinctions of species all over the planet^1,2,6,7,10,33,37–39,43^. Our observation of extinctions over successive and independent time periods strongly reflects the magnitude of post-industrial impacts as drivers of multisystemic global environmental degradation, with species loss not only increasing drastically in the last ∼200 years^6,10^ (see results), but also vastly expanding across continental lands. The projections of future extinctions we provide reinforce the critical nature of this ongoing biodiversity crisis. Our findings that post-industrial extinction patterns are roughly similar among mammals, birds and reptiles strongly contrast with the explosive rates of amphibian extinctions over the last century. This singularity in the extinction trends of amphibians globally has been suggested to be aggravated by the chytridiomycosis fungal disease – deemed nature’s most destructive wildlife pathogen^39,72,75^.

Humans have been a driver of biodiversity loss for thousands of years^61,76^. However, our findings align with the wider evidence that the magnitude of environmental degradation caused by industrialisation, and the growth of human population that has come with it, rapidly triggered a catastrophic acceleration of species extinctions from across the tree of life. Therefore, the aggregated impacts resulting from a progressively growing human population are likely to continue to aggravate the loss of biodiversity during the 21^st^ century. In fact, even at the lowest confidence interval, the UN predicts a net population increase of nearly 1 billion people globally between 2030 and 2100, which leads to an estimated loss of 576 species during that period in a best-case scenario (compared to the estimated loss of an additional 1,244 species in a worst-case scenario where net human population increases in >4 billion people). Collectively, our study is a further warning to humanity about the urgent need to implement cooperative actions (e.g., the adoption of the UN Sustainable Development Goals) and to stabilise the world’s population (‘within a framework that ensures social integrity’^69^) to mitigate the impacts of climate change on the annihilation of life on Earth and on the wellbeing of humans globally.

## Materials and methods

### Tetrapod extinction data

We collated a global-scale database covering all species of tetrapod vertebrates (amphibians, reptiles, birds and mammals) currently declared extinct (EX) or possibly extinct (PE) by the 2021 IUCN Red List^23^. PE species are formally a subcategory within critically endangered (CR) species that are thought to be extinct, but which require confirmation. The latest recorded observations of PE species date back as far as the 1820s, with multiple other species last seen only during the 19^th^ century^6,23^ (Supplementary Table S1). Our analyses exclude species classed as extinct in the wild (EW) given that many of these species currently kept in captivity are the subject of breeding programs and thus, their recurrence in the wild is not necessarily irreversible. For each of the 540 tetrapod species classed as EX and PE (‘extinct’ hereafter) we searched for the year of their last recorded sighting in the wild. This information was not available for 31 species given lack of records, the existence of fossil remains only, or because records in videos or pictures suggests that they may still occur in the wild (Supplementary Table S1), and thus, they were excluded from the analyses. In other cases, the year since last observation is not exact, with data available for a specific decade (e.g., 1920s) or even century (e.g., 1500-1600). In these cases, we assigned the species a ‘conservative’ estimated date set to be the last year of that time period, e.g., ‘1929s’ and ‘1599s’, respectively (Supplementary Table S1).

### Temporal trajectories of species extinctions

To reconstruct the temporal trajectories of extinctions (Fig. 1), years since extinction (year of last record in the wild) were organised into time-lapses of two decades for each tetrapod class separately, from 1460 to 2000 (where 1464 is our earliest recorded extinction). Whereas extinctions occurred from the year 2000 were included in our analyses, they were treated separately from reconstructions of the main temporal trajectories given that categorisation of species as EX or PE require, among other strict criteria, ‘exhaustive surveys’ across the entire known historical range of the species ‘over a timeframe appropriate to its life cycle and life form’^77^ (e.g., after over a century since last observation, some species remain PE). The consistently lower number of EX+PE species recorded post-2000 (relative to the class-specific tendencies recorded for the past few decades) is, therefore, importantly due to application of criteria rather than to a reduction in extinction events.

To perform quantitative temporal analyses of extinction patterns, all species with non-exact extinction years and those that went extinct after the year 2000 were removed from the analysis, leaving a total of 415 species. Pearson’s Chi-squared tests were used to test our null hypothesis that the mean numbers of extinctions before and after the industrial revolution (set at 1760 for these tests) are the same by comparing the expected means (equal for both groups) and observed means. These analyses were done for total extinctions, continent-only extinctions, and island-only extinctions (see next section for details about species’ continental and insular distributions). A second Chi-squared test was performed on amphibians only, comparing extinctions between 1951-1975 and 1976-2000 over 5-year intervals.

Mann-Kendall trend tests^78^ were performed on data from pre-1760 and post-1760 separately to determine trend changes in numbers of extinctions (in 20-year intervals) before and after the industrial revolution, split by class and island/continent. The strength of the trend is indicated by Kendall’s tau (*τ*), which is a value between -1 and 1, where a negative value indicates a negative trend and the larger the value, the stronger the trend. A 2-sided *p*-value is used to quantify statistical significance. Note, there were no recorded amphibian or reptile extinctions pre-1760 and no recorded bird extinctions on the continent pre-1760 with exact extinction dates.

### Spatial patterns of species extinctions

Species distribution maps (shapefiles) were accessed through The IUCN Red List of Threatened Species^23^. For species with no data available on IUCN, distribution data were reconstructed from the literature (Supplementary Table S4). To establish biogeographic patterns of extinctions through time, species were treated separately as island or mainland endemics (Supplementary Table S1). Given the vast variation in the territorial surface of non-continental landmasses, we treated all territories with a surface of 150,000km^2^ or less as islands, whereas all remaining oceanic territories (e.g., Madagascar, Australia, Papua New Guinea) were classed as mainland. Species that occurred both on mainland and islands (Supplementary Table S1) were classed within the mainland group as the logic behind this classification aimed to capture the impacts that island isolation exerts on species that face anthropogenic threats.

To calculate the number of species per grid-cell at different time lapses, we merged multipolygon and point data for each separated group, by first buffering the point data to 0.00001 degree (∼3.42±0.4 m^2^) and then binding both layers. Subsequently, we used a world grid of 1×1 degree cell-size and joined the species distribution data at four temporal ranges: 1800-1849, 1850-1899, 1900-1949, and 1950-1999, to count the number of species per grid-cell. The same was done for all groups together, yet this time considering five temporal ranges corresponding to the centuries: 1400s+1500s, 1600s, 1700s, 1800s, and 1900s.

To improve the visualization (Fig. 2), we calculated and plotted the centroids of the grid cells. The scale colour is the same for all maps, from blue (1-2 species), green (3-6 species) and red (7-9 species). All spatial analyses were done in the R software^79^ using the ‘sf’ package^80^.

### Extinction predictions based on human population growth scenarios

Future tetrapod extinctions as a function of human population count were predicted using human population growth projections from the United Nations^81^, and accounted for median, upper 95% confidence interval and lower 95% confidence interval growth estimates. The predictions are based on 5-year intervals, reflecting a balance between number of points and statistical power.

Extinctions occurring after the year 2000 were not included due to the time required to adequately assess the extinction of species (see above), and therefore, the years 2005, 2010, 2015 and 2020 were estimated from the observed HPG-extinctions pre-2000 using our chosen model. For past human populations, we used data from the Our World in Data resource (https://ourworldindata.org/grapher/population-past-future), which provides 5-year intervals from the year 1800. Therefore, only extinctions after 1800 were used in this analysis.

At the stage of initial data exploration, a strong correlation was found between human population growth and number of extinctions (*r*(38)=0.763, *P*<0.001), providing evidence for the validity of model creation. Over-dispersion in the data was also detected. As such, Generalised Linear Models (GLMs) of the quasipoisson family were generated using both the log link and identity link functions, and negative binomials GLMs were generated using 5 repeats of 5-fold cross-validation in R package ‘caret’^82^, using the R package ‘MASS’^83^. Two outliers were highlighted in diagnostic plots and thus, models were run both with and without these outliers. Models were compared using root mean squared error (RMSE), and a quasipoisson GLM with identity link function run on data from which outliers were removed was chosen as the final model. Although RMSE were very similar for this and the negative binomial GLM with identity link function on the data with outliers removed, the quasipoisson outperformed the negative binomial when the model was run on the full dataset (Supplementary Figure S1). All analyses were performed in R version 4.1.0^79^.

## Acknowledgments

The completion of this project has been funded by a grant from the School of Biological Sciences at Queen’s University Belfast to DPD. LPH is grateful to Nottingham Trent University for full PhD funding. CF is funded by a NERC PhD DTP research program. JVJ and LEBG are funded by PhD scholarships from the Department of Economy of Northern Ireland.

