## Supplementary Table S1 for "Temporal and spatial patterns of vertebrate extinctions during the Anthropocene"

**Supplementary Table S1.** Dataset used in this study. Data have been taken from the IUCN Red List database. Extinction statuses are abbreviated as EX (species declared as confirmed extinctions) and PE(CR) (species declared possibly extinct). Year refers to the year when the species was last recorded in the wild. In cases where the year is denoted with an ‘S’ indicate that the record is imprecise and thus, the species has been conservatively assigned the latest year of the decade (e.g., 1959s for “1950s”) or century (e.g., 1699s for “1600s”) when the species was last seen (see methods in the main text for details).

| **Class** | **Order** | **Genus** | **Species** | **Geographic_Region** | **Distribution** | **IUCN** | **Year** |
| --- | --- | --- | --- | --- | --- | --- | --- |
| Amphibia  Amphibia  Amphibia  Amphibia  Amphibia  Amphibia  Amphibia  Amphibia  Amphibia  Amphibia  Amphibia  Amphibia  Amphibia  Amphibia  Amphibia  Amphibia  Amphibia  Amphibia  Amphibia  Amphibia  Amphibia  Amphibia  Amphibia  Amphibia  Amphibia  Amphibia  Amphibia  Amphibia  Amphibia  Amphibia  Amphibia  Amphibia  Amphibia  Amphibia  Amphibia  Amphibia  Amphibia  Amphibia  Amphibia  Amphibia  Amphibia  Amphibia  Amphibia  Amphibia  Amphibia  Amphibia  Amphibia  Amphibia  Amphibia  Amphibia  Amphibia  Amphibia  Amphibia  Amphibia  Amphibia  Amphibia  Amphibia  Amphibia  Amphibia  Amphibia  Amphibia  Amphibia  Amphibia  Amphibia  Amphibia  Amphibia  Amphibia  Amphibia  Amphibia  Amphibia  Amphibia  Amphibia  Amphibia  Amphibia  Amphibia  Amphibia  Amphibia  Amphibia  Amphibia  Amphibia  Amphibia  Amphibia  Amphibia  Amphibia  Amphibia  Amphibia  Amphibia  Amphibia  Amphibia  Amphibia  Amphibia  Amphibia  Amphibia  Amphibia  Amphibia  Amphibia  Amphibia  Amphibia  Amphibia  Amphibia  Amphibia  Amphibia  Amphibia  Amphibia  Amphibia  Amphibia  Amphibia  Amphibia  Amphibia  Amphibia  Amphibia  Amphibia  Amphibia  Amphibia  Amphibia  Amphibia  Amphibia  Amphibia  Amphibia  Amphibia  Amphibia  Amphibia  Amphibia  Amphibia  Amphibia  Amphibia  Amphibia  Amphibia  Amphibia  Amphibia  Amphibia  Amphibia  Amphibia  Amphibia  Amphibia  Amphibia  Amphibia  Amphibia  Amphibia  Amphibia  Amphibia  Amphibia  Amphibia  Amphibia  Amphibia  Amphibia  Amphibia  Amphibia  Amphibia  Amphibia  Amphibia  Amphibia  Amphibia  Amphibia  Amphibia  Amphibia  Amphibia  Amphibia  Amphibia  Amphibia  Amphibia  Amphibia  Amphibia  Amphibia  Amphibia  Amphibia  Amphibia  Amphibia  Amphibia  Amphibia  Amphibia  Reptilia  Reptilia  Reptilia  Reptilia  Reptilia  Reptilia  Reptilia  Reptilia  Reptilia  Reptilia  Reptilia  Reptilia  Reptilia  Reptilia  Reptilia  Reptilia  Reptilia  Reptilia  Reptilia  Reptilia  Reptilia  Reptilia  Reptilia  Reptilia  Reptilia  Reptilia  Reptilia  Reptilia  Reptilia  Reptilia  Reptilia  Reptilia  Reptilia  Reptilia  Reptilia  Reptilia  Reptilia  Reptilia  Reptilia  Reptilia  Reptilia  Reptilia  Reptilia  Reptilia  Reptilia  Reptilia  Reptilia  Reptilia  Reptilia  Reptilia  Reptilia  Reptilia  Reptilia  Reptilia  Reptilia  Reptilia  Reptilia  Reptilia  Reptilia  Reptilia  Reptilia  Reptilia  Reptilia  Reptilia  Reptilia  Reptilia  Aves  Aves  Aves  Aves  Aves  Aves  Aves  Aves  Aves  Aves  Aves  Aves  Aves  Aves  Aves  Aves  Aves  Aves  Aves  Aves  Aves  Aves  Aves  Aves  Aves  Aves  Aves  Aves  Aves  Aves  Aves  Aves  Aves  Aves  Aves  Aves  Aves  Aves  Aves  Aves  Aves  Aves  Aves  Aves  Aves  Aves  Aves  Aves  Aves  Aves  Aves  Aves  Aves  Aves  Aves  Aves  Aves  Aves  Aves  Aves  Aves  Aves  Aves  Aves  Aves  Aves  Aves  Aves  Aves  Aves  Aves  Aves  Aves  Aves  Aves  Aves  Aves  Aves  Aves  Aves  Aves  Aves  Aves  Aves  Aves  Aves  Aves  Aves  Aves  Aves  Aves  Aves  Aves  Aves  Aves  Aves  Aves  Aves  Aves  Aves  Aves  Aves  Aves  Aves  Aves  Aves  Aves  Aves  Aves  Aves  Aves  Aves  Aves  Aves  Aves  Aves  Aves  Aves  Aves  Aves  Aves  Aves  Aves  Aves  Aves  Aves  Aves  Aves  Aves  Aves  Aves  Aves  Aves  Aves  Aves  Aves  Aves  Aves  Aves  Aves  Aves  Aves  Aves  Aves  Aves  Aves  Aves  Aves  Aves  Aves  Aves  Aves  Aves  Aves  Aves  Aves  Aves  Aves  Aves  Aves  Aves  Aves  Aves  Aves  Aves  Aves  Aves  Aves  Aves  Aves  Aves  Aves  Aves  Aves  Aves  Aves  Aves  Aves  Aves  Aves  Aves  Mammalia  Mammalia  Mammalia  Mammalia  Mammalia  Mammalia  Mammalia  Mammalia  Mammalia  Mammalia  Mammalia  Mammalia  Mammalia  Mammalia  Mammalia  Mammalia  Mammalia  Mammalia  Mammalia  Mammalia  Mammalia  Mammalia  Mammalia  Mammalia  Mammalia  Mammalia  Mammalia  Mammalia  Mammalia  Mammalia  Mammalia  Mammalia  Mammalia  Mammalia  Mammalia  Mammalia  Mammalia  Mammalia  Mammalia  Mammalia  Mammalia  Mammalia  Mammalia  Mammalia  Mammalia  Mammalia  Mammalia  Mammalia  Mammalia  Mammalia  Mammalia  Mammalia  Mammalia  Mammalia  Mammalia  Mammalia  Mammalia  Mammalia  Mammalia  Mammalia  Mammalia  Mammalia  Mammalia  Mammalia  Mammalia  Mammalia  Mammalia  Mammalia  Mammalia  Mammalia  Mammalia  Mammalia  Mammalia  Mammalia  Mammalia  Mammalia  Mammalia  Mammalia  Mammalia  Mammalia  Mammalia  Mammalia  Mammalia  Mammalia  Mammalia  Mammalia  Mammalia  Mammalia  Mammalia  Mammalia | Anura  Anura  Anura  Anura  Anura  Anura  Anura  Anura  Anura  Anura  Anura  Anura  Anura  Anura  Anura  Anura  Anura  Anura  Anura  Anura  Anura  Anura  Anura  Anura  Anura  Anura  Anura  Anura  Anura  Anura  Anura  Anura  Anura  Anura  Anura  Anura  Anura  Anura  Anura  Anura  Anura  Anura  Anura  Anura  Anura  Anura  Anura  Anura  Anura  Anura  Anura  Anura  Anura  Anura  Anura  Anura  Anura  Anura  Anura  Anura  Anura  Anura  Anura  Anura  Anura  Anura  Anura  Anura  Anura  Anura  Anura  Anura  Anura  Anura  Anura  Anura  Anura  Anura  Anura  Anura  Anura  Anura  Anura  Anura  Anura  Anura  Anura  Anura  Anura  Anura  Anura  Anura  Anura  Anura  Anura  Anura  Anura  Anura  Anura  Anura  Anura  Anura  Anura  Anura  Anura  Anura  Anura  Anura  Anura  Anura  Anura  Anura  Anura  Anura  Anura  Anura  Anura  Anura  Anura  Anura  Anura  Anura  Anura  Anura  Anura  Anura  Anura  Anura  Anura  Anura  Anura  Anura  Anura  Anura  Anura  Anura  Anura  Anura  Anura  Anura  Anura  Anura  Anura  Anura  Anura  Anura  Anura  Anura  Anura  Anura  Anura  Anura  Anura  Anura  Anura  Anura  Anura  Anura  Anura  Caudata  Caudata  Caudata  Caudata  Caudata  Caudata  Caudata  Caudata  Caudata  Caudata  Caudata  Caudata  Squamata  Squamata  Squamata  Squamata  Squamata  Squamata  Squamata  Squamata  Squamata  Squamata  Squamata  Squamata  Squamata  Squamata  Squamata  Squamata  Squamata  Squamata  Squamata  Squamata  Squamata  Squamata  Squamata  Squamata  Squamata  Squamata  Squamata  Squamata  Squamata  Squamata  Squamata  Squamata  Squamata  Squamata  Squamata  Squamata  Squamata  Squamata  Squamata  Squamata  Squamata  Squamata  Squamata  Squamata  Squamata_Snake  Squamata_Snake  Squamata_Snake  Squamata_Snake  Squamata_Snake  Squamata_Snake  Squamata_Snake  Squamata_Snake  Squamata_Snake  Squamata_Snake  Squamata_Snake  Squamata_Snake  Squamata_Snake  Testudines  Testudines  Testudines  Testudines  Testudines  Testudines  Testudines  Testudines  Testudines  Accipitriformes  Anseriformes  Anseriformes  Anseriformes  Anseriformes  Anseriformes  Anseriformes  Anseriformes  Bucerotiformes  Caprimulgiformes  Caprimulgiformes  Caprimulgiformes  Caprimulgiformes  Caprimulgiformes  Charadriiformes  Charadriiformes  Charadriiformes  Charadriiformes  Charadriiformes  Charadriiformes  Charadriiformes  Charadriiformes  Charadriiformes  Charadriiformes  Columbiformes  Columbiformes  Columbiformes  Columbiformes  Columbiformes  Columbiformes  Columbiformes  Columbiformes  Columbiformes  Columbiformes  Columbiformes  Columbiformes  Columbiformes  Columbiformes  Columbiformes  Columbiformes  Cuculiformes  Cuculiformes  Falconiformes  Falconiformes  Galliformes  Gruiformes  Gruiformes  Gruiformes  Gruiformes  Gruiformes  Gruiformes  Gruiformes  Gruiformes  Gruiformes  Gruiformes  Gruiformes  Gruiformes  Gruiformes  Gruiformes  Gruiformes  Gruiformes  Gruiformes  Gruiformes  Gruiformes  Gruiformes  Gruiformes  Gruiformes  Gruiformes  Gruiformes  Passeriformes  Passeriformes  Passeriformes  Passeriformes  Passeriformes  Passeriformes  Passeriformes  Passeriformes  Passeriformes  Passeriformes  Passeriformes  Passeriformes  Passeriformes  Passeriformes  Passeriformes  Passeriformes  Passeriformes  Passeriformes  Passeriformes  Passeriformes  Passeriformes  Passeriformes  Passeriformes  Passeriformes  Passeriformes  Passeriformes  Passeriformes  Passeriformes  Passeriformes  Passeriformes  Passeriformes  Passeriformes  Passeriformes  Passeriformes  Passeriformes  Passeriformes  Passeriformes  Passeriformes  Passeriformes  Passeriformes  Passeriformes  Passeriformes  Passeriformes  Passeriformes  Passeriformes  Passeriformes  Passeriformes  Passeriformes  Passeriformes  Passeriformes  Passeriformes  Passeriformes  Passeriformes  Passeriformes  Passeriformes  Passeriformes  Passeriformes  Passeriformes  Passeriformes  Passeriformes  Passeriformes  Passeriformes  Passeriformes  Passeriformes  Passeriformes  Passeriformes  Passeriformes  Passeriformes  Passeriformes  Pelecaniformes  Pelecaniformes  Pelecaniformes  Pelecaniformes  Pelecaniformes  Pelecaniformes  Piciformes  Piciformes  Podicipediformes  Podicipediformes  Podicipediformes  Procellariiformes  Procellariiformes  Procellariiformes  Procellariiformes  Psittaciformes  Psittaciformes  Psittaciformes  Psittaciformes  Psittaciformes  Psittaciformes  Psittaciformes  Psittaciformes  Psittaciformes  Psittaciformes  Psittaciformes  Psittaciformes  Psittaciformes  Psittaciformes  Psittaciformes  Psittaciformes  Psittaciformes  Psittaciformes  Psittaciformes  Strigiformes  Strigiformes  Strigiformes  Strigiformes  Strigiformes  Strigiformes  Struthioniformes  Struthioniformes  Suliformes  Afrosoricida  Carnivora  Carnivora  Carnivora  Carnivora  Carnivora  Carnivora  Cetartiodactyla  Cetartiodactyla  Cetartiodactyla  Cetartiodactyla  Cetartiodactyla  Cetartiodactyla  Cetartiodactyla  Cetartiodactyla  Chiroptera  Chiroptera  Chiroptera  Chiroptera  Chiroptera  Chiroptera  Chiroptera  Chiroptera  Chiroptera  Chiroptera  Chiroptera  Chiroptera  Chiroptera  Dasyuromorphia  Didelphimorphia  Didelphimorphia  Diprotodontia  Diprotodontia  Diprotodontia  Diprotodontia  Diprotodontia  Diprotodontia  Diprotodontia  Diprotodontia  Diprotodontia  Eulipotyphla  Lagomorpha  Peramelemorphia  Peramelemorphia  Peramelemorphia  Primates  Primates  Primates  Primates  Rodentia  Rodentia  Rodentia  Rodentia  Rodentia  Rodentia  Rodentia  Rodentia  Rodentia  Rodentia  Rodentia  Rodentia  Rodentia  Rodentia  Rodentia  Rodentia  Rodentia  Rodentia  Rodentia  Rodentia  Rodentia  Rodentia  Rodentia  Rodentia  Rodentia  Rodentia  Rodentia  Rodentia  Rodentia  Rodentia  Rodentia  Rodentia  Rodentia  Rodentia  Rodentia  Rodentia  Rodentia  Rodentia  Rodentia  Rodentia  Sirenia | *"Prostherapis"*  *Allobates*  *Altiphrynoides*  *Andinobates*  *Andinobates*  *Aromobates*  *Aromobates*  *Aromobates*  *Aromobates*  *Aromobates*  *Arthroleptides*  *Arthroleptis*  *Arthroleptis*  *Atelopus*  *Atelopus*  *Atelopus*  *Atelopus*  *Atelopus*  *Atelopus*  *Atelopus*  *Atelopus*  *Atelopus*  *Atelopus*  *Atelopus*  *Atelopus*  *Atelopus*  *Atelopus*  *Atelopus*  *Atelopus*  *Atelopus*  *Atelopus*  *Atelopus*  *Atelopus*  *Atelopus*  *Atelopus*  *Atelopus*  *Atelopus*  *Atelopus*  *Atelopus*  *Atelopus*  *Atelopus*  *Atelopus*  *Atelopus*  *Atelopus*  *Atelopus*  *Atelopus*  *Atelopus*  *Atelopus*  *Atelopus*  *Atelopus*  *Centrolene*  *Centrolene*  *Craugastor*  *Craugastor*  *Craugastor*  *Craugastor*  *Craugastor*  *Craugastor*  *Craugastor*  *Craugastor*  *Craugastor*  *Craugastor*  *Craugastor*  *Craugastor*  *Craugastor*  *Craugastor*  *Craugastor*  *Craugastor*  *Dryophytes*  *Ecnomiohyla*  *Ecnomiohyla*  *Ectopoglossus*  *Eleutherodactylus*  *Eleutherodactylus*  *Eleutherodactylus*  *Eleutherodactylus*  *Eleutherodactylus*  *Eleutherodactylus*  *Gastrotheca*  *Gastrotheca*  *Holoaden*  *Hyloscirtus*  *Hyloxalus*  *Hyloxalus*  *Hyloxalus*  *Incilius*  *Incilius*  *Incilius*  *Isthmohyla*  *Lithobates*  *Lithobates*  *Lithobates*  *Litoria*  *Litoria*  *Melanophryniscus*  *Nannophryne*  *Nannophrys*  *Nectophrynoides*  *Nymphargus*  *Oophaga*  *Parhoplophryne*  *Paruwrobates*  *Peltophryne*  *Philautus*  *Phrynobatrachus*  *Phrynobatrachus*  *Phrynomedusa*  *Plectrohyla*  *Pristimantis*  *Pristimantis*  *Pristimantis*  *Pristimantis*  *Pristimantis*  *Proceratophrys*  *Pseudophilautus*  *Pseudophilautus*  *Pseudophilautus*  *Pseudophilautus*  *Pseudophilautus*  *Pseudophilautus*  *Pseudophilautus*  *Pseudophilautus*  *Pseudophilautus*  *Pseudophilautus*  *Pseudophilautus*  *Pseudophilautus*  *Pseudophilautus*  *Pseudophilautus*  *Pseudophilautus*  *Pseudophilautus*  *Pseudophilautus*  *Rana*  *Rheobatrachus*  *Rheobatrachus*  *Rhinella*  *Rhinella*  *Rhinoderma*  *Sarcohyla*  *Sarcohyla*  *Sarcohyla*  *Sarcohyla*  *Sarcohyla*  *Sarcohyla*  *Sarcohyla*  *Scutiger*  *Strabomantis*  *Strabomantis*  *Taudactylus*  *Telmatobius*  *Telmatobius*  *Telmatobius*  *Telmatobius*  *Telmatobius*  *Telmatobius*  *Telmatobius*  *Telmatobius*  *Telmatobius*  *Telmatobius*  *Telmatobius*  *Aquiloeurycea*  *Bolitoglossa*  *Cynops*  *Oedipina*  *Plethodon*  *Pseudoeurycea*  *Pseudoeurycea*  *Pseudoeurycea*  *Pseudoeurycea*  *Pseudoeurycea*  *Pseudoeurycea*  *Thorius*  *Alinea*  *Alinea*  *Anolis*  *Capitellum*  *Capitellum*  *Capitellum*  *Celestus*  *Celestus*  *Chioninia*  *Contomastix*  *Copeoglossum*  *Cyclura*  *Cynisca*  *Emoia*  *Holcosus*  *Hoplodactylus*  *Leiocephalus*  *Leiocephalus*  *Leiocephalus*  *Leiocephalus*  *Leiocephalus*  *Leiocephalus*  *Leiolopisma*  *Lepidoblepharis*  *Liolaemus*  *Mabuya*  *Mabuya*  *Mabuya*  *Phelsuma*  *Pholidoscelis*  *Scelotes*  *Sphaerodactylus*  *Sphaerodactylus*  *Sphaerodactylus*  *Spondylurus*  *Spondylurus*  *Spondylurus*  *Spondylurus*  *Spondylurus*  *Spondylurus*  *Spondylurus*  *Stenocercus*  *Tachygyia*  *Tetradactylus*  *Anilios*  *Bolyeria*  *Borikenophis*  *Calamaria*  *Clelia*  *Erythrolamprus*  *Erythrolamprus*  *Hypsirhynchus*  *Madatyphlops*  *Mitophis*  *Omoadiphas*  *Pseudoxyrhopus*  *Trimetopon*  *Chelonoidis*  *Chelonoidis*  *Chelonoidis*  *Cylindraspis*  *Cylindraspis*  *Cylindraspis*  *Cylindraspis*  *Cylindraspis*  *Pelusios*  *Bermuteo*  *Alopochen*  *Alopochen*  *Anas*  *Anas*  *Camptorhynchus*  *Chenonetta*  *Mergus*  *Upupa*  *Chlorostilbon*  *Chlorostilbon*  *Eriocnemis*  *Eurostopodus*  *Siphonorhis*  *Coenocorypha*  *Coenocorypha*  *Haematopus*  *Numenius*  *Pinguinus*  *Prosobonia*  *Prosobonia*  *Prosobonia*  *Turnix*  *Vanellus*  *Alectroenas*  *Alectroenas*  *Alopecoenas*  *Alopecoenas*  *Caloenas*  *Columba*  *Columba*  *Columba*  *Ectopistes*  *Microgoura*  *Nesoenas*  *Nesoenas*  *Nesoenas*  *Pezophaps*  *Ptilinopus*  *Raphus*  *Coua*  *Nannococcyx*  *Caracara*  *Falco*  *Coturnix*  *Aphanapteryx*  *Cabalus*  *Diaphorapteryx*  *Dryolimnas*  *Erythromachus*  *Fulica*  *Gallinula*  *Hypotaenidia*  *Hypotaenidia*  *Hypotaenidia*  *Hypotaenidia*  *Laterallus*  *Mundia*  *Porphyrio*  *Porphyrio*  *Porphyrio*  *Porphyrio*  *Porphyrio*  *Tribonyx*  *Zapornia*  *Zapornia*  *Zapornia*  *Zapornia*  *Zapornia*  *Acrocephalus*  *Acrocephalus*  *Acrocephalus*  *Acrocephalus*  *Acrocephalus*  *Akialoa*  *Akialoa*  *Akialoa*  *Akialoa*  *Anthornis*  *Aplonis*  *Aplonis*  *Aplonis*  *Aplonis*  *Callaeas*  *Carpodacus*  *Chaetoptila*  *Chloridops*  *Cichlocolaptes*  *Ciridops*  *Drepanis*  *Drepanis*  *Dysmorodrepanis*  *Foudia*  *Fregilupus*  *Gerygone*  *Hemignathus*  *Hemignathus*  *Hemignathus*  *Heteralocha*  *Himatione*  *Loxops*  *Loxops*  *Melamprosops*  *Moho*  *Moho*  *Moho*  *Moho*  *Myadestes*  *Myadestes*  *Myadestes*  *Myiagra*  *Necropsar*  *Nesillas*  *Paroreomyza*  *Paroreomyza*  *Philydor*  *Pipilo*  *Pomarea*  *Pomarea*  *Pomarea*  *Pomarea*  *Poodytes*  *Psittirostra*  *Pyrocephalus*  *Quiscalus*  *Rhodacanthis*  *Rhodacanthis*  *Traversia*  *Turdus*  *Turnagra*  *Turnagra*  *Vermivora*  *Viridonia*  *Xenicus*  *Zoothera*  *Zosterops*  *Zosterops*  *Zosterops*  *Ixobrychus*  *Nyctanassa*  *Nycticorax*  *Nycticorax*  *Nycticorax*  *Threskiornis*  *Campephilus*  *Colaptes*  *Podiceps*  *Podilymbus*  *Tachybaptus*  *Bulweria*  *Hydrobates*  *Pterodroma*  *Pterodroma*  *Alexandrinus*  *Amazona*  *Amazona*  *Anodorhynchus*  *Ara*  *Charmosyna*  *Conuropsis*  *Cyanoramphus*  *Cyanoramphus*  *Eclectus*  *Lophopsittacus*  *Lophopsittacus*  *Mascarinus*  *Necropsittacus*  *Nestor*  *Palaeornis*  *Psephotellus*  *Psittacara*  *Pyrrhura*  *Aegolius*  *Glaucidium*  *Mascarenotus*  *Mascarenotus*  *Mascarenotus*  *Sceloglaux*  *Dromaius*  *Dromaius*  *Urile*  *Cryptochloris*  *Cryptoprocta*  *Dusicyon*  *Dusicyon*  *Neomonachus*  *Neovison*  *Zalophus*  *Bos*  *Bos*  *Gazella*  *Gazella*  *Hippotragus*  *Lipotes*  *Rucervus*  *Sus*  *Murina*  *Mystacina*  *Nyctophilus*  *Pipistrellus*  *Pipistrellus*  *Pteralopex*  *Pteropus*  *Pteropus*  *Pteropus*  *Pteropus*  *Pteropus*  *Pteropus*  *Pteropus*  *Thylacinus*  *Cryptonanus*  *Monodelphis*  *Bettongia*  *Caloprymnus*  *Dendrolagus*  *Lagorchestes*  *Lagorchestes*  *Macropus*  *Onychogalea*  *Phalanger*  *Potorous*  *Crocidura*  *Prolagus*  *Chaeropus*  *Macrotis*  *Perameles*  *Nycticebus*  *Palaeopropithecus*  *Piliocolobus*  *Xenothrix*  *Conilurus*  *Dipodomys*  *Geocapromys*  *Gyldenstolpia*  *Isolobodon*  *Juscelinomys*  *Lagostomus*  *Leporillus*  *Megalomys*  *Megalomys*  *Melanomys*  *Melomys*  *Mesocapromys*  *Mesocapromys*  *Nesoryzomys*  *Nesoryzomys*  *Nilopegamys*  *Noronhomys*  *Notomys*  *Notomys*  *Notomys*  *Notomys*  *Oligoryzomys*  *Oryzomys*  *Oryzomys*  *Pennatomys*  *Peromyscus*  *Peromyscus*  *Peromyscus*  *Pseudomys*  *Pseudomys*  *Pseudomys*  *Rattus*  *Rattus*  *Spalax*  *Tylomys*  *Tylomys*  *Uromys*  *Uromys*  *Uromys*  *Hydrodamalis* | *dunni*  *ranoides*  *osgoodi*  *abditus*  *viridis*  *alboguttatus*  *haydeeae*  *leopardalis*  *nocturnus*  *serranus*  *dutoiti*  *kutogundua*  *troglodytes*  *angelito*  *ardila*  *carbonerensis*  *chiriquiensis*  *chocoensis*  *ebenoides*  *erythropus*  *eusebiodiazi*  *farci*  *guanujo*  *halihelos*  *longirostris*  *lynchi*  *mindoensis*  *minutulus*  *monohernandezii*  *nicefori*  *onorei*  *orcesi*  *oxyrhynchus*  *pachydermus*  *pastuso*  *peruensis*  *petersi*  *petriruizi*  *pictiventris*  *pinangoi*  *planispina*  *podocarpus*  *quimbaya*  *senex*  *sernai*  *simulatus*  *sonsonensis*  *sorianoi*  *subornatus*  *vogli*  *geckoideum*  *pipilatum*  *adamastus*  *anciano*  *andi*  *catalinae*  *chrysozetetes*  *cruzi*  *epochthidius*  *fecundus*  *merendonensis*  *myllomyllon*  *olanchano*  *omoaensis*  *phasma*  *punctariolus*  *rhyacobatrachus*  *trachydermus*  *bocourti*  *echinata*  *rabborum*  *atopoglossus*  *eneidae*  *glanduliferoides*  *karlschmidti*  *orcutti*  *schmidti*  *semipalmatus*  *angustifrons*  *antomia*  *bradei*  *chlorosteus*  *abditaurantius*  *edwardsi*  *ruizi*  *fastidiosus*  *majordomus*  *periglenes*  *calypsa*  *fisheri*  *pueblae*  *tlaloci*  *castanea*  *piperata*  *peritus*  *cophotis*  *guentheri*  *poyntoni*  *truebae*  *speciosa*  *usambarica*  *andinus*  *fluviatica*  *jacobsoni*  *manengoubensis*  *njiomock*  *fimbriata*  *pycnochila*  *albericoi*  *anotis*  *bernali*  *molybrignus*  *phragmipleuron*  *moratoi*  *adspersus*  *dimbullae*  *eximius*  *extirpo*  *halyi*  *leucorhinus*  *maia*  *malcolmsmithi*  *nanus*  *nasutus*  *oxyrhynchus*  *pardus*  *rugatus*  *temporalis*  *variabilis*  *zal*  *zimmeri*  *chevronta*  *silus*  *vitellinus*  *chrysophora*  *rostrata*  *rufum*  *calvicollina*  *charadricola*  *cyanomma*  *pachyderma*  *psarosema*  *sabrina*  *siopela*  *maculatus*  *cadenai*  *necerus*  *diurnus*  *bolivianus*  *ceiorum*  *cirrhacelis*  *edaphonastes*  *espadai*  *laticeps*  *mendelsoni*  *niger*  *pefauri*  *sibiricus*  *vellardi*  *praecellens*  *nussbaumi*  *wolterstorffi*  *petiola*  *ainsworthi*  *anitae*  *aquatica*  *brunnata*  *exspectata*  *teotepec*  *unguidentis*  *longicaudus*  *lanceolata*  *luciae*  *roosevelti*  *mariagalantae*  *metallicum*  *parvicruzae*  *anelpistus*  *occiduus*  *coctei*  *charrua*  *redondae*  *onchiopsis*  *gansi*  *nativitatis*  *orcesi*  *delcourti*  *cuneus*  *endomychus*  *eremitus*  *herminieri*  *pratensis*  *rhutidira*  *mauritiana*  *miyatai*  *cranwelli*  *hispaniolae*  *mabouya*  *montserratae*  *gigas*  *cineraceus*  *guentheri*  *elasmorhynchus*  *lazelli*  *williamsi*  *anegadae*  *haitiae*  *lineolatus*  *magnacruzae*  *martinae*  *monitae*  *spilonotus*  *haenschi*  *microlepis*  *eastwoodae*  *insperatus*  *multocarinata*  *sanctaecrucis*  *prakkei*  *errabunda*  *cursor*  *perfuscus*  *melanichnus*  *cariei*  *leptepileptus*  *cannula*  *ankafinaensis*  *viquezi*  *abingdonii*  *niger*  *phantasticus*  *indica*  *inepta*  *peltastes*  *triserrata*  *vosmaeri*  *seychellensis*  *avivorus*  *kervazoi*  *mauritiana*  *marecula*  *theodori*  *labradorius*  *finschi*  *australis*  *antaios*  *bracei*  *elegans*  *godini*  *exul*  *americana*  *barrierensis*  *iredalei*  *meadewaldoi*  *borealis*  *impennis*  *cancellata*  *ellisi*  *leucoptera*  *novaecaledoniae*  *macropterus*  *nitidissimus*  *payandeei*  *ferrugineus*  *salamonis*  *maculata*  *jouyi*  *thiriouxi*  *versicolor*  *migratorius*  *meeki*  *cicur*  *duboisi*  *rodericanus*  *solitaria*  *mercierii*  *cucullatus*  *delalandei*  *psix*  *lutosa*  *duboisi*  *novaezelandiae*  *bonasia*  *modestus*  *hawkinsi*  *augusti*  *leguati*  *newtonii*  *nesiotis*  *dieffenbachii*  *pacifica*  *poeciloptera*  *wakensis*  *podarces*  *elpenor*  *albus*  *caerulescens*  *kukwiedei*  *mantelli*  *paepae*  *hodgenorum*  *astrictocarpus*  *monasa*  *nigra*  *palmeri*  *sandwichensis*  *astrolabii*  *luscinius*  *musae*  *nijoi*  *yamashinae*  *ellisiana*  *lanaiensis*  *obscura*  *stejnegeri*  *melanocephala*  *corvina*  *fusca*  *mavornata*  *ulietensis*  *cinereus*  *ferreorostris*  *angustipluma*  *kona*  *mazarbarnetti*  *anna*  *funerea*  *pacifica*  *munroi*  *delloni*  *varius*  *insularis*  *affinis*  *hanapepe*  *lucidus*  *acutirostris*  *fraithii*  *ochraceus*  *wolstenholmei*  *phaeosoma*  *apicalis*  *bishopi*  *braccatus*  *nobilis*  *lanaiensis*  *myadestinus*  *woahensis*  *freycineti*  *rodericanus*  *aldabrana*  *flammea*  *maculata*  *novaesi*  *naufragus*  *fluxa*  *mira*  *nukuhivae*  *pomarea*  *rufescens*  *psittacea*  *dubius*  *palustris*  *flaviceps*  *palmeri*  *lyalli*  *ravidus*  *capensis*  *tanagra*  *bachmanii*  *sagittirostris*  *longipes*  *terrestris*  *conspicillatus*  *semiflavus*  *strenuus*  *novaezelandiae*  *carcinocatactes*  *duboisi*  *mauritianus*  *megacephalus*  *solitarius*  *imperialis*  *oceanicus*  *andinus*  *gigas*  *rufolavatus*  *bifax*  *macrodactylus*  *caribbaea*  *rupinarum*  *exsul*  *martinicana*  *violacea*  *glaucus*  *tricolor*  *diadema*  *carolinensis*  *ulietanus*  *zealandicus*  *infectus*  *bensoni*  *mauritianus*  *mascarin*  *rodricanus*  *productus*  *wardi*  *pulcherrimus*  *labati*  *subandina*  *gradyi*  *mooreorum*  *grucheti*  *murivorus*  *sauzieri*  *albifacies*  *baudinianus*  *minor*  *perspicillatus*  *wintoni*  *spelea*  *australis*  *avus*  *tropicalis*  *macrodon*  *japonicus*  *primigenius*  *sauveli*  *bilkis*  *saudiya*  *leucophaeus*  *vexillifer*  *schomburgki*  *bucculentus*  *tenebrosa*  *robusta*  *howensis*  *murrayi*  *sturdeei*  *pulchra*  *allenorum*  *aruensis*  *brunneus*  *coxi*  *pilosus*  *subniger*  *tokudae*  *cynocephalus*  *ignitus*  *unistriata*  *anhydra*  *campestris*  *mayri*  *asomatus*  *leporides*  *greyi*  *lunata*  *matanim*  *platyops*  *trichura*  *sardus*  *ecaudatus*  *leucura*  *eremiana*  *bancanus*  *ingens*  *waldroni*  *mcgregori*  *albipes*  *gravipes*  *thoracatus*  *fronto*  *portoricensis*  *candango*  *crassus*  *apicalis*  *desmarestii*  *luciae*  *zunigae*  *rubicola*  *nanus*  *sanfelipensis*  *darwini*  *indefessus*  *plumbeus*  *vespuccii*  *amplus*  *longicaudatus*  *macrotis*  *mordax*  *victus*  *antillarum*  *nelsoni*  *nivalis*  *guardia*  *mekisturus*  *pembertoni*  *auritus*  *glaucus*  *gouldii*  *macleari*  *nativitatis*  *istricus*  *bullaris*  *tumbalensis*  *emmae*  *imperator*  *porculus*  *gigas* | South America  South America  Africa  South America  South America  South America  South America  South America  South America  South America  Africa  Africa  Africa  South America  South America  South America  South America  South America  South America  South America  South America  South America  South America  South America  South America  South America  South America  South America  South America  South America  South America  South America  South America  South America  South America  South America  South America  South America  South America  South America  South America  South America  South America  South America  South America  South America  South America  South America  South America  South America  South America  South America  Central America  Central America  Central America  Central America  Central America  Central America  Central America  Central America  Central America  Central America  Central America  Central America  Central America  Central America  Central America  Central America  Central America  North America  Central America  South America  Puerto Rico  Hispaniola  Puerto Rico  Jamaica  Hispaniola  Hispaniola  South America  South America  South America  South America  South America  South America  South America  South America  South America  South America  Central America  North America  North America  North America  Australia  Australia  South America  South America  Sri Lanka  Africa  South America  Central America  Africa  South America  Hispaniola  Java  Africa  Africa  South America  Central America  South America  South America  South America  South America  South America  South America  Sri Lanka  Sri Lanka  Sri Lanka  Sri Lanka  Sri Lanka  Sri Lanka  Sri Lanka  Sri Lanka  Sri Lanka  Sri Lanka  Sri Lanka  Sri Lanka  Sri Lanka  Sri Lanka  Sri Lanka  Sri Lanka  Sri Lanka  Asia  Australia  Australia  Central America  South America  South America  Central America  Central America  Central America  Central America  Central America  Central America  Central America  Asia  South America  South America  Australia  South America  South America  South America  South America  South America  South America  South America  South America  South America  South America  South America  North America  Central America  Asia  Central America  North America  North America  North America  North/Central America  Central America  North America  North America  North America  Barbados, Caribbean  Saint Lucia, Caribbean  Puerto Rican Bank Islands, Caribbean  Guadeloupe  Martinique  St. Croix in the US Virgin Islands, Caribbean  Hispaniola, Caribbean  Jamaica  Cape Verde islands  Uruguay  Redonda Island, Antigua and Barbuda  Navassa Island  Nigeria  Christmas Island  Ecuador  New Zealand  Anguilla, Barbuda, Antigua, and Guadeloupe  Hispaniola, Caribbean  Navassa Island  Martinique  Hispaniola, Caribbean  Hispaniola, Caribbean  Mauritius  Colombia  Bolivia  Hispaniola, Caribbean  Martinique, Caribbean  Montserrat Island, Caribbean  Rodrigues, Mauritius  Grand Ilet off Petit-Bourg, Caribbean  South Africa  Hispaniola, Caribbean  Hispaniola, Caribbean  Hispaniola, Caribbean  Anegada, Caribbean  Haiti  Hispaniola, Caribbean  St. Croix and Green Cay, US Virgin Islands, Caribbean  St. Martin, Caribbean  islet Monito, Puerto Rico, Caribbean  St. Thomas and St. John, US Virgin Islands, Caribbean  Ecuador  Tonga Island  Africa  Australia  Round Island, Mauritius  St. Croix, U.S. Virgin Islands, Caribbean  Malaysia  Saint Lucia  Martinique and Rocher du Diamant, Caribbean  Barbados  Hispaniola, Caribbean  Mauritius  Hispaniola, Caribbean  Honduras  Madagascar  Costa Rica  Pinta, Galapagos  Floreana, Galapagos  Fernandina Island, Galapagos  Reunion  Mauritius  Rodriguez  Mauritius  Rodriguez  Seychelles  Bermuda  Reunion  Mauritius  Amsterdam Island, French Southern Territories  Mauritius  North America  North and South Islands, NZ  Auckland Islands, NZ  St Helena  Bahamas  Jamaica or the north Bahamas  Ecuador  New Caledonia  Jamaica  Little Barrier Island, NZ  Stewart Island, NZ  Canary Islands, Spain  Americas  North America and Europe  Christmas Island, Kiribati  Moorea, in the Society Islands, French Polynesia  Society Islands, French Polynesia  New Caledonia  Java, Indonesia  Mauritius  Rodrigues, Mauritius  Tanna Island, Vanuatu  Makira and Ramos, Solomon Islands  Unknown  Nansei Shoto Islands, Japan  Mauritius  Bonin Islands, Japan  North America  Choiseul, Solomon Islands  Mauritius  Reunion  Rodrigues, Mauritius  Rodrigues, Mauritius  Marquesas, French Polynesia  Mauritius  Ile de Sainte-Marie, Madagascar  St Helena  Guadalupe Island, Mexico  Reunion  North, South and Great Barrier Islands, NZ  Mauritius  Chatham, Mangere and Pitt Islands, NZ  Chatham Islands, NZ  Reunion  Rodrigues, Mauritius  Reunion/Mauritius  Tristan da Cunha, UK  Chatham, Mangere and Pitt Islands, NZ  Tahiti, French Polynesia  Fiji Islands  Wake Island, US Minor Outlying Islands  St Helena  Ascension Island  Lord Howe Island, Australia  Reunion  New Caledonia  North Island, NZ  Marquesas Islands, French Polynesia  North and South Islands, NZ  St Helena  Kosrae, Caroline Islands, Micronesia  Tahiti, French Polynesia  Laysan Island, Hawaii  Hawaii  Gambier Islands, French Polynesia  Guam  Society Islands, French Polynesia  Aguijan, Northern Mariana Islands  Pagan, Northern Mariana Islands  O'ahu, Hawaii  Lanai, Hawaii  Hawaii  Kauai, Hawaii  Chatham, Mangere and Little Mangere Islands, NZ  Kosrae, Caroline Islands, Micronesia  Norfolk and Lord Howe Island, Australia  Mauke, Cook Islands  Raiatea, French Polynesia  South and Stewart Islands, NZ  Peel Island, Bonin, Japan  Hawaii  Hawaii  Brazil  Hawaii  Molokai, Hawaii  Hawaii  Lanai, Hawaii  Reunion  Reunion  Lord Howe Island, Australia  Maui, Hawaiian Islands  Kauai, Hawaii  O'ahu, Hawaii  North Island, NZ  Laysan Island, Hawaii  Maui, Hawaii  O'ahu, Hawaii  Maui, Hawaii  O'ahu, Hawaii  Hawaiian Islands  Kauai, Hawaii  Hawaii  Maui, Lanai and Molokai Islands, Hawaii  Kauai, Hawaii  O'ahu, in Hawaii  Guam  Rodrigues, Mauritius  Ile Malabar, Aldabra, Seychelles  Molokai, Hawaii  O'ahu, Hawaii  Brazil  Bermuda  Marquesas Islands, French Polynesia  Island of Ua Pou, Marquesas Islands, French Polynesia  Nuku Hiva, Marquesas Islands, French Polynesia  Maupiti, Society Islands, French Polynesia  Chatham Islands, NZ  Hawaiian Islands  San Cristóbal, Galapagos  Mexico  Kona, Hawaii  Hawaii  Stephens Island, NZ  Grand Cayman, Cayman Islands  South Island, NZ  North Island, NZ  North America and Caribbean  Hawaii  South, Stewart and North Islands, NZ  Peel Island, Bonin, Japan  Guam  Marianne Island, Seychelles  Lord Howe Island, Australia  South and North Islands, NZ  Bermuda  Reunion  Mauritius  Rodrigues, Mauritius  Reunion/Rodrigues  Mexico  Bermuda  Colombia  Guatemala  Madagascar  St Helena  Guadalupe, Mexico  Jamaica  St Helena  Rodrigues, Mauritius  Martinique  Guadeloupe  South America  Cuba  New Caledonia  North America  Raiatea, French Polynesia  Tahiti, French Polynesia  Tonga  Mauritius  Mauritius  Reunion  Rodrigues, Mauritius  Norfolk Island and Phillip Island, Australia  Mahé and Silhouette, Seychelles  Australia  Guadeloupe  Colombia  Bermuda  Brazil  Reunion  Rodrigues, Mauritius  Mauritius  South, Stewart and North Islands, NZ  Kangaroo Island, Australia  King Island, Australia  Russia/Berling Island  South Africa  Madagascar  Falkland/Malvinas Islands  South America  Caribbean Sea  North America  Northwest Pacific  Eurasia and Africa  South East Asia  Yemen  Arabian Peninsula  South Africa  China  Thailand  South East Asia  Tsushima Island, Japan  Big South Cape Island  Lord Howe Island, Australia  Christmas Island, Australia  Hahajima Island, Bonin, Japan  Guadacanal, Solomon Islands  Upolu, Samoa  Aru Islands, Indonesia  Percy Islands, Australia  Samoan Archipelago  Palau  Mauritius and Réunion  Guam  Tasmania  Argentina  South America  Australia  Australia  Papua New Guinea  Australia  Australia  Australia  Australia  Papua New Guinea  Australia  Christmas Island, Australia  Corsica, Sardinia and other Mediterranean islands  Australia  Australia  Australia  Bangka Island, Indonesia  Madagascar  Africa  Jamaica  Australia  Mexico  Little Swan Island, Honduras  Argentina  Hispaniola  Brazil  Peru  Australia  Martinique, Caribbean  St. Lucia  Peru  Bramble Cay, Torres Strait, Australia  Cuba  Cayos de San Felipe Island, Caribbean  Santa Cruz Island, Galapagos  Santa Cruz and Baltra Island, Galapagos  Africa  Fernando de Noronha Island, Brazil  Australia  Australia  Australia  Australia  Saint Vincent, Caribbean  Jamaica  María Madre Island, Mexico  Saint Kitts Bank, Caribbean  Ángel de la Guarda Island, Mexico  Mexico  San Pedro Nolasco Island, Mexico  Kangaroo Island and Australia  Australia  Australia  Christmas Island, Australia  Christmas Island, Australia  Romania  Mexico  Mexico  Paidaido Islands, Indonesia  Guadalcanal, Solomon Islands  Guadalcanal, Solomon Islands  Bering Sea | Continent  Continent  Continent  Continent  Continent  Continent  Continent  Continent  Continent  Continent  Continent  Continent  Continent  Continent  Continent  Continent  Continent  Continent  Continent  Continent  Continent  Continent  Continent  Continent  Continent  Continent  Continent  Continent  Continent  Continent  Continent  Continent  Continent  Continent  Continent  Continent  Continent  Continent  Continent  Continent  Continent  Continent  Continent  Continent  Continent  Continent  Continent  Continent  Continent  Continent  Continent  Continent  Continent  Continent  Continent  Continent  Continent  Continent  Continent  Continent  Continent  Continent  Continent  Continent  Continent  Continent  Continent  Continent  Continent  Continent  Continent  Continent  Island  Island  Island  Island  Island  Island  Continent  Continent  Continent  Continent  Continent  Continent  Continent  Continent  Continent  Continent  Continent  Continent  Continent  Continent  Continent  Continent  Continent  Continent  Island  Continent  Continent  Continent  Continent  Continent  Island  Island  Continent  Continent  Continent  Continent  Continent  Continent  Continent  Continent  Continent  Continent  Island  Island  Island  Island  Island  Island  Island  Island  Island  Island  Island  Island  Island  Island  Island  Island  Island  Continent  Continent  Continent  Continent  Continent  Continent  Continent  Continent  Continent  Continent  Continent  Continent  Continent  Continent  Continent  Continent  Continent  Continent  Continent  Continent  Continent  Continent  Continent  Continent  Continent  Continent  Continent  Continent  Continent  Continent  Continent  Continent  Continent  Continent  Continent  Continent  Continent  Continent  Continent  Continent  Island  Island  Island  Island  Island  Island  Island  Island  Island  Continent  Island  Island  Continent  Island  Continent  Island  Island  Island  Island  Island  Island  Island  Island  Continent  Continent  Island  Island  Island  Island  Island  Continent  Island  Island  Island  Island  Island  Island  Island  Island  Island  Island  Continent  Island  Continent  Continent  Island  Island  Continent  Island  Island  Island  Island  Island  Island  Continent  Continent  Continent  Island  Island  Island  Island  Island  Island  Island  Island  Island  Island  Island  Island  Island  Island  Continent  Island  Island  Island  Island  Island  Continent  Island  Island  Island  Island  Island  Continent  Continent/Island  Island  Island  Island  Island  Island  Island  Island  Island  Island  Unknown  Island  Island  Island  Continent  Island  Island  Island  Island  Island  Island  Island  Island  Island  Island  Island  Island  Island  Island  Island  Island  Island  Island  Island  Island  Island  Island  Island  Island  Island  Island  Island  Island  Island  Island  Continent  Island  Island  Island  Island  Island  Island  Island  Island  Island  Island  Island  Island  Island  Island  Island  Island  Island  Island  Island  Island  Island  Island  Island  Continent  Island  Island  Island  Island  Island  Island  Island  Island  Island  Island  Island  Island  Island  Island  Island  Island  Island  Island  Island  Island  Island  Island  Island  Island  Island  Island  Island  Continent  Island  Island  Island  Island  Island  Island  Island  Island  Continent  Island  Island  Island  Island  Island  Island  Continent/Island  Island  Island  Island  Island  Island  Island  Island  Island  Island  Island  Island  Island  Continent  Island  Continent  Continent  Continent  Island  Island  Island  Island  Island  Island  Island  Continent  Island  Island  Continent  Island  Island  Island  Island  Island  Island  Island  Island  Island  Continent  Island  Continent  Island  Continent  Island  Island  Island  Island  Island  Island  Continent/Island  Continent  Continent  Island  Continent  NA  Continent  NA  Continent  Continent  Continent  Continent  Continent  Continent  Continent  Continent  Island  Island  Island  Island  Island  Island  Island  Island  Island  Island  Island  Island  Island  Island  Continent  Continent  Continent  Continent  Continent  Continent  Continent  Continent  Continent  Continent  Continent  Island  Island  Continent  Continent  Continent  Island  Continent  Continent  Island  Continent  Continent  Island  Continent  Island  Continent  Continent  Continent  Island  Island  Continent  Island  Island  Island  Island  Island  Continent  Island  Continent  Continent  Continent  Continent  Island  Island  Island  Island  Island  Continent  Island  Continent/Island  Continent  Continent  Island  Island  Continent  Continent  Continent  Island  Island  Island  NA | PE(CR)  PE(CR)  PE(CR)  PE(CR)  PE(CR)  PE(CR)  PE(CR)  PE(CR)  PE(CR)  PE(CR)  PE(CR)  PE(CR)  PE(CR)  PE(CR)  PE(CR)  PE(CR)  EX  PE(CR)  PE(CR)  PE(CR)  PE(CR)  PE(CR)  PE(CR)  PE(CR)  EX  PE(CR)  PE(CR)  PE(CR)  PE(CR)  PE(CR)  PE(CR)  PE(CR)  PE(CR)  PE(CR)  PE(CR)  PE(CR)  PE(CR)  PE(CR)  PE(CR)  PE(CR)  PE(CR)  PE(CR)  PE(CR)  EX  PE(CR)  PE(CR)  PE(CR)  PE(CR)  PE(CR)  EX  PE(CR)  PE(CR)  PE(CR)  EX  PE(CR)  PE(CR)  PE(CR)  PE(CR)  PE(CR)  PE(CR)  PE(CR)  EX  PE(CR)  EX  PE(CR)  PE(CR)  PE(CR)  PE(CR)  PE(CR)  PE(CR)  PE(CR)  PE(CR)  PE(CR)  PE(CR)  PE(CR)  PE(CR)  PE(CR)  PE(CR)  PE(CR)  PE(CR)  PE(CR)  PE(CR)  PE(CR)  PE(CR)  PE(CR)  PE(CR)  PE(CR)  EX  PE(CR)  EX  PE(CR)  PE(CR)  PE(CR)  PE(CR)  PE(CR)  PE(CR)  EX  PE(CR)  PE(CR)  EX  PE(CR)  PE(CR)  PE(CR)  PE(CR)  PE(CR)  PE(CR)  EX  PE(CR)  PE(CR)  PE(CR)  PE(CR)  PE(CR)  PE(CR)  PE(CR)  EX  EX  EX  EX  EX  EX  EX  EX  EX  EX  EX  EX  EX  EX  EX  EX  EX  PE(CR)  EX  EX  PE(CR)  PE(CR)  PE(CR)  PE(CR)  PE(CR)  PE(CR)  PE(CR)  PE(CR)  PE(CR)  PE(CR)  PE(CR)  PE(CR)  PE(CR)  EX  PE(CR)  PE(CR)  PE(CR)  PE(CR)  PE(CR)  PE(CR)  PE(CR)  PE(CR)  PE(CR)  PE(CR)  PE(CR)  PE(CR)  PE(CR)  EX  PE(CR)  EX  PE(CR)  PE(CR)  PE(CR)  EX  PE(CR)  PE(CR)  PE(CR)  PE(CR)  EX  PE(CR)  PE(CR)  PE(CR)  PE(CR)  PE(CR)  PE(CR)  EX  EX  EX  EX  PE(CR)  EX  PE(CR)  EX  EX  PE(CR)  EX  EX  PE(CR)  PE(CR)  EX  PE(CR)  PE(CR)  PE(CR)  PE(CR)  PE(CR)  EX  EX  EX  PE(CR)  PE(CR)  PE(CR)  PE(CR)  PE(CR)  PE(CR)  PE(CR)  PE(CR)  PE(CR)  PE(CR)  PE(CR)  EX  EX  PE(CR)  EX  PE(CR)  PE(CR)  EX  PE(CR)  EX  PE(CR)  EX  PE(CR)  PE(CR)  PE(CR)  PE(CR)  EX  EX  PE(CR)  EX  EX  EX  EX  EX  EX  EX  EX  EX  EX  EX  EX  EX  EX  EX  EX  EX  PE(CR)  PE(CR)  PE(CR)  EX  EX  EX  PE(CR)  EX  EX  EX  EX  PE(CR)  PE(CR)  EX  EX  EX  EX  EX  EX  EX  EX  EX  EX  EX  EX  EX  EX  EX  EX  EX  EX  EX  EX  EX  EX  EX  EX  EX  EX  EX  EX  EX  EX  EX  EX  EX  EX  EX  EX  EX  EX  EX  EX  EX  EX  EX  EX  EX  EX  EX  EX  EX  EX  EX  EX  EX  EX  EX  EX  EX  EX  EX  PE(CR)  EX  EX  EX  EX  EX  EX  EX  EX  EX  EX  EX  PE(CR)  PE(CR)  EX  EX  EX  PE(CR)  EX  EX  EX  EX  EX  EX  PE(CR)  EX  EX  EX  EX  EX  EX  PE(CR)  EX  EX  EX  PE(CR)  EX  EX  EX  PE(CR)  EX  EX  EX  EX  EX  EX  EX  EX  PE(CR)  EX  EX  EX  EX  EX  EX  EX  EX  EX  EX  EX  EX  PE(CR)  EX  EX  EX  EX  EX  PE(CR)  PE(CR)  EX  EX  EX  EX  PE(CR)  EX  PE(CR)  EX  EX  EX  EX  EX  EX  EX  EX  EX  EX  EX  EX  PE(CR)  EX  PE(CR)  EX  EX  EX  EX  EX  EX  EX  PE(CR)  EX  EX  EX  EX  EX  EX  EX  PE(CR)  EX  EX  EX  PE(CR)  EX  EX  PE(CR)  PE(CR)  EX  EX  EX  PE(CR)  EX  PE(CR)  EX  EX  EX  EX  EX  EX  EX  PE(CR)  EX  EX  PE(CR)  EX  EX  EX  EX  PE(CR)  EX  PE(CR)  EX  EX  EX  EX  PE(CR)  EX  PE(CR)  EX  EX  PE(CR)  EX  PE(CR)  EX  EX  EX  EX  EX  EX  PE(CR)  EX  PE(CR)  PE(CR)  EX  EX  PE(CR)  EX  EX  EX  EX  EX  EX  EX  EX  EX  PE(CR)  PE(CR)  EX  EX  EX  EX  EX  EX  PE(CR)  PE(CR)  PE(CR)  PE(CR)  PE(CR)  PE(CR)  EX | 1977  2001  2003  2004  2005  1995  1999S  1995  1991  1999S  1962  1930  1962  2000  1989  1998  1996  1998  2019s  2004  1997  2003  1988  1988  1989  1984  1989  1985  1982  1988  1990  1988  1994  1995  1993  1992  1996  1998  1996  1997  1985  1994  1992  1986  2001  2003  1996  1990  1993  1957  1997  1977  1983  1990  1990  1990  1989  1980  1999  1989  1968  1978  1999S  1974  1992  2010  1982  1989  1984  1962  2008  1999S  1990  1985  1988  1989S  1989S  1985  1999S  1997  1976  1979  2000  1996  1979  1990  1980  1989  2002  1942  1948  1985  1980  1973  1953  2005  1882  2003  2005  1992  1929s  1999S  1981  1912  2009S  2010  1929s  1954  1986  1974  1981  1999S  1983  1990  1886  1940  1933  1882  1899  1859s  1876  1927  1869  1869  1872  1859s  1927  1864  1858  1949s  1927  1983  1981  1985  1996  1914  1981  1980  1961  1980  1940  1978  1970  1979S  1979S  1982  1995  1979  2007  1984  1981  1999  1998  1993  2007  1994  1976  2003  1987  1940  1998  1979  1995  1964  1976  1978  1989S  1976  1964  1976  1998  1889  1889  1932  1839s  1829S  1875  1977  1859s  1912  1977  1873  1878  1969s  2010  1959  1869  1699s  1976  1868  1837  1966  1978  1699s  1964  1973  1937  1889  1984  1899s  1914  1887  1966  1960  1978  1966  1858  1985  2000  1969s  1993  1877  1901  1839  1928  1992  1975  1899  1893  1844  1968  1963  1910  1699s  1984  1989  1889s  1927  1972  1859s  1906  1800  1749s  1800  1749s  1800  1895  1603  1672  1698  1800  1710  1875  1599s  1902  1550  1900  1900  1850  1939  1860  1870  1964  1950  1963  1852  1850  1800  1800  1911  1939  1840  1690s  1800  1927  1928  1936  1730  1889  1914  1904  1730  1750  1799s  1770  1950  1662  1834  1800  1900  1672  1875  1700  1900  1895  1670  1726  1693  1900  1872  1940  1973  1945  1550  1815  1834  1730  1860  1850  1937  1699s  1550  1875  1800  1944  1884  1839  1969  1900  1995  1969  1837  1892  1940  1969  1906  1880  1923  1825  1812  2007  1900  1900  1894  2007  1937  1907  1898  1920  1672  1860  1936  1896  1899  1899  1907  1923  1988  1930  2004  1850  1981  1987  1934  1980  1989  1825  1983  1726  1983  1963  1985  2011  1612  1982  2010  1951  1850  1900  1989  1987  1910  1891  1896  1900  1938  1963  1955  1988  1901  1972  1889  1983  1892  1928  1900  1610  1674  1700  1761  1750  1956  1623  1977  1986  1985  1550  1912  1879  1550  1875  1800  1800  2001  1850  1913  1918  1773  1850  1799s  1800  1700  1834  1761  1851  1900  1927  1800  1949  1699s  2001  1650  1726  1859  1970  1827  1850  1850  1937  1658  1876  1464  1952  1860  1951  1627  1970  1951  1970  1800  2002  1932  1892  1962  1967  1972  2009  1889  1991  1856  1992  1890s  1841  1874  1859  1968  1933  1962  1899  1933  1935  1928  1959s  1890  1979s  1959s  1997  1875  1985  1799s  1959s  1969s  1943  1937  1620  2008  1799s  1862  1986  1959s  1896  1525  1960  1910  1979s  1897  1881  1949  2009  1951  1978  1939s  1934  1920s  1500s  1896  1902  1844  1840s  1892  1877  1897  1930  1991  1957  1931  1850  1899s  1857  1904  1898  1983  1970  1970  1946  1888  1888  1768 |
