## Supplementary Table S2 for "Temporal and spatial patterns of vertebrate extinctions during the Anthropocene"

**Supplementary Table S2.** Mann-Kendall trend test results measuring temporal trend changes in 20-year intervals for all tetrapod taxa globally, and split by both class and island/continent. Tests were not performed on amphibian and reptile extinctions or continental bird extinctions pre-1760 since there are no extinction records available with exact extinction dates. Kendall’s τ is a measure of trend strength, valued between -1 and 1, the significance of which is quantified by a 2-sided p-value. Significant p-values are highlighted in bold italic.

| **Model** | **Taxa** | **Kendall’s** τ | **2-sided *P*-value** |
| --- | --- | --- | --- |
| **Pre-industrial era (1460-1760)**  Global  Global  Global  Island  Island  Island  Continent  Continent  **Post-industrial era (1760-2000)**  Global  Global  Global  Global  Global  Island  Island  Island  Island  Island  Continent  Continent  Continent  Continent  Continent | Tetrapods  Aves  Mammalia  Tetrapods  Aves  Mammalia  Tetrapods  Mammalia  Tetrapods  Amphibia  Aves  Mammalia  Reptilia  Tetrapods  Amphibia  Aves  Mammalia  Reptilia  Tetrapods  Amphibia  Aves  Mammalia  Reptilia | 0.4700  0.5649  -0.1354  0.5469  0.5649  -0.1649  -0.0527  -0.0527  0.7613  0.8161  0.4416  0.5338  0.6672  0.5128  0.4199  0.4212  0.5854  0.5767  0.8466  0.8006  0.5947  0.6319  0.7621 | ***0.0183***  ***0.0051***  0.5711  ***0.0065***  ***0.0051***  0.5152  0.8557  0.8557  ***0.0004***  ***0.0003***  ***0.0433***  ***0.0159***  ***0.0025***  ***0.0173***  0.0794  0.0564  ***0.0095***  ***0.0105***  ***0.0001***  ***0.0006***  ***0.0109***  ***0.0054***  ***0.0012*** |
