## Supplementary Table S3 for "Temporal and spatial patterns of vertebrate extinctions during the Anthropocene"

**Supplementary Table S3.** Numbers of tetrapod species predicted to have become extinct from the year 2000 (and their 95% confidence intervals) as a function of sustained increases in human population density, based on the actual relationships between species gone extinct or potentially extinct and human population growth between the years 1800-2000. The predicted numbers of extinct species have been approximated while the CI have not. See main text for details.

| Year | Human Population Density | Number of extinct species | Lower 95% CI | Upper 95% CI |
| --- | --- | --- | --- | --- |
| 1805  1810  1815  1820  1825  1830  1835  1840  1845  1850  1855  1860  1865  1870  1875  1880  1885  1890  1895  1900  1905  1910  1915  1920  1925  1930  1935  1940  1945  1950  1955  1960  1965  1970  1975  1980  1985  1990  1995  2000  2005  2010  2015  2020  2025  2030  2035  2040  2045  2050  2055  2060  2065  2070  2075  2080  2085  2090  2095  2100 | 1,001,758,044  1,022,630,197  1,054,586,106  1,092,947,422  1,128,342,960  1,166,619,859  1,194,777,021  1,224,118,692  1,250,936,428  1,278,698,170  1,289,344,086  1,303,155,263  1,322,223,328  1,348,154,272  1,383,207,445  1,425,998,653  1,477,744,363  1,536,461,987  1,588,397,643  1,647,405,022  1,716,473,051  1,793,323,592  1,855,949,027  1,926,217,425  2,008,052,088  2,104,067,378  2,210,412,972  2,327,357,739  2,417,413,700  2,536,605,808  2,773,019,915  3,035,160,180  3,339,583,509  3,700,685,676  4,079,480,473  4,458,274,952  4,870,921,665  5,327,529,078  5,744,212,929  6,143,776,621  6,542,205,330  6,957,137,521  7,380,117,870  7,794,799,000  8,184,437,000  8,548,487,000  8,887,524,000  9,198,847,000  9,481,803,000  9,735,034,000  9,958,099,000  10,151,470,000  10,317,879,000  10,459,240,000  10,577,288,000  10,673,904,000  10,750,662,000  10,809,892,000  10,851,860,000  10,875,394,000 | 0  0  2  0  2  1  3  5  4  12  3  16  4  3  9  8  4  10  8  7  10  7  13  2  2  14  9  15  4  6  13  7  18  20  10  30  36  38  27  21  35  38  40  43  45  47  49  51  52  54  55  56  57  58  59  59  60  60  60  61 | 5  11  26.52941012  28.25424877  30.01120403  31.73263338  33.34932681  34.85928233  36.26507198  37.55563756  38.72838237  39.77776353  40.70202033  41.50315966  42.19253795  42.77811142  43.26708693  43.66727026  43.98519116  44.23050728  44.40432519  44.50179426 | 8  16  43.77927322  46.90808934  50.09893188  53.22822052  56.16930395  58.91781054  61.47789432  63.82901862  65.9661417  67.87892129  69.56395951  71.02477068  72.28195583  73.34994771  74.24183454  74.97181349  75.55176659  75.99929133  76.31639263  76.49421169 |
