## Supplementary Figure S1 for "Temporal and spatial patterns of vertebrate extinctions during the Anthropocene"

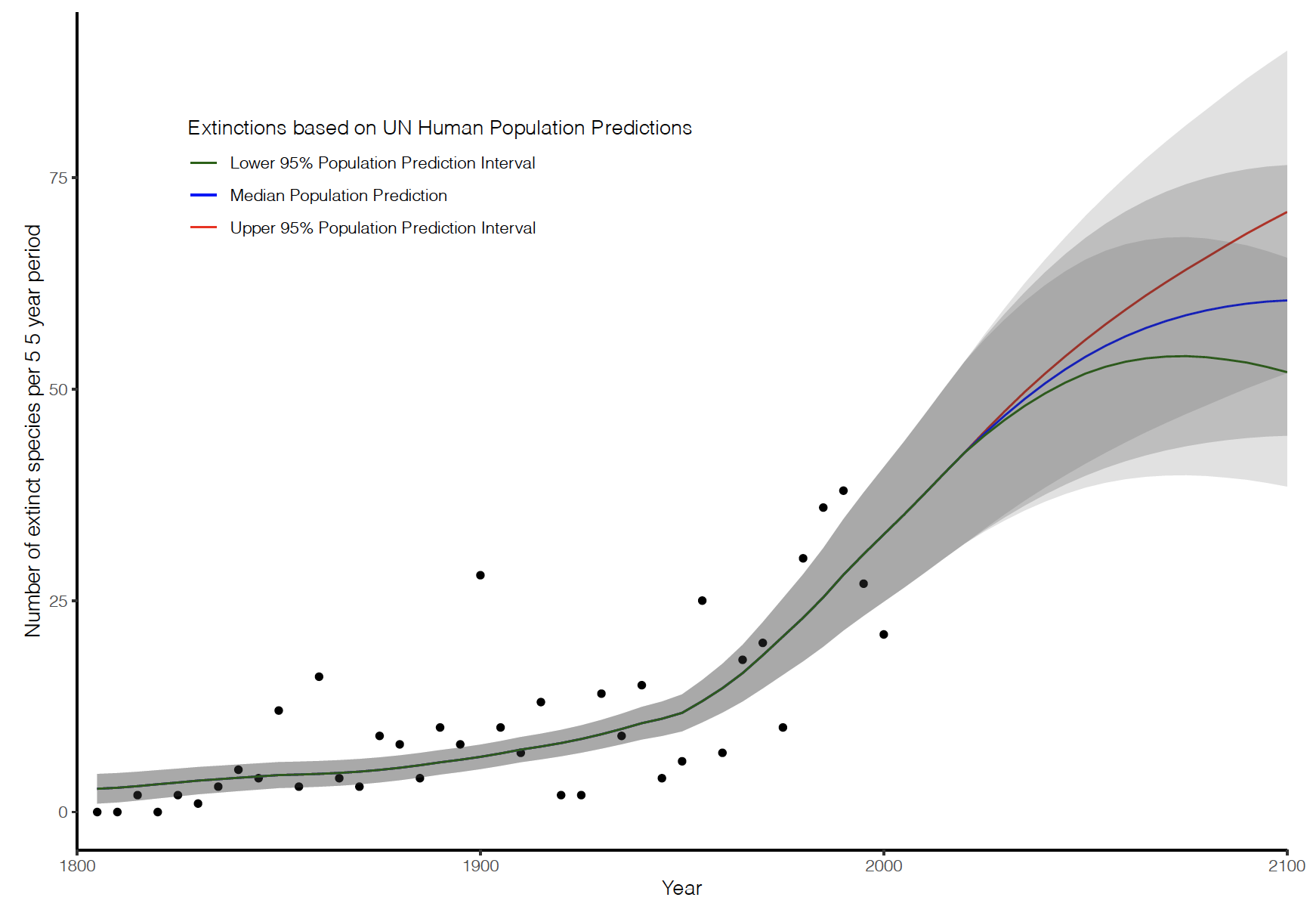

**Supplementary Figure S1.** Number of recorded extinctions in 5-year intervals from 1801-2000 with predictions of number of extinct species in 5-year intervals between 2001 and 2100 based on human population growth scenarios from the UN *with outliers included*. Green indicates the lowest bound of human population growth prediction, blue the median and red the upper bound. The numbers of extinctions are raw figures that indicate the number of species that are lost for every five-year period, with the year of each datapoint signifying the final year in the period. 95% confidence intervals are shown in grey.
